# Recent social experience alters song behavior in *Drosophila*

**DOI:** 10.1101/2025.09.04.674308

**Authors:** Frederic A. Roemschied, Elise C Ireland, Adam J Calhoun, Minseung Choi, Osama M. Ahmed, Mala Murthy

## Abstract

Innate behaviors are hardwired into the nervous system, allowing animals to produce them instinctively and without prior learning. This includes social behaviors (e.g., aggression, courtship) in most animals. If and how learning shapes such innate behaviors has not been systematically explored. Here, we investigate learning from social feedback in *Drosophila*. We use closed-loop optogenetics to systematically perturb the social experience of individual flies during courtship, and find that altered feedback changes the male fly’s innate courtship strategy, opening the door for investigating the neural and molecular basis of social plasticity in flies.

**Teaser:** Male flies adjust their innate communication strategy in response to systematic perturbation of social experience.

## Introduction

Acoustic communication is an important behavior across the animal kingdom, used to attract potential mates over long distances, or to warn conspecifics about dangers. Acoustic signal production is learned in only a small number of species (1). For the rest, the behavior is thought to be hard-wired and to not require learning. Typically, males produce sounds as part of their courtship ritual, and they are born knowing how to do so. This is the case for *Drosophila melanogaster* males, who, as part of their innate courtship behavior, generate complex wing-born acoustic signals (2, 3). Male song is organized into sequences consisting of alternations between three syllables (two ‘pulse’ modes and one ‘sine’ mode (4)). While male flies do not learn their songs, their singing behavior is nonetheless highly flexible (5). Males select each song syllable as well as song amplitude dependent on dynamically changing sensory feedback from females (6, 7) and relative to changing internal states (8). Given this flexibility, we asked whether male flies can learn to alter the context in which they produce song. This type of learning has been referred to as usage learning (9, 10). There have been limited studies of social experience-dependent learning in male *Drosophila*; these studies typically measure the suppression of courtship via associative learning ((11, 12, 13), but see (14, 15) for examples of male mate choice learning and mate copying, respectively), but not whether the strategy males use to court (for example, singing sine song only when near the female (5)) can be adjusted. Such usage learning is important because it allows for adaptability in optimizing innate behaviors for novel circumstances. Usage learning has been explored in other model systems, including birds, seals, monkeys, and killer whales (16, 17, 18). Here, we use real-time, closed-loop optogenetics to perturb female feedback during courtship song production, and show that *Drosophila melanogaster* males are capable of usage learning. We find that males that experience perturbed female feedback alter the context in which they sing with new females, opening the door to dissecting the genetic, neural circuit, and evolutionary mechanisms of such learning.

## Results

### A Novel Assay for Perturbing Male Experience During Singing

During courtship in *Drosophila*, females accelerate to elicit male song (6) and slow down in response to song (19, 20) - her movements are a major cue used by the male to shape both chasing and singing (21, 22). To explore usage learning in *Drosophila*, we reasoned that targeting female motion cues would provide a salient teaching signal to the courting male. Although backward walking is a natural behavior, females almost never walk backward during courtship. We developed an assay to trigger backward walking in females via optogenetic activation (23) of ‘moonwalker’ descending neurons (MDNs (24)) triggered on male pulse song production (**Fig. 1A-D**). Despite the natural entanglement of pulse and sine song modes in bouts (5), our perturbation was specific to pulse song (**Fig. S1A-D**). Pulse song is produced both when far and near the female, whereas sine song predominantly occurs when near a female (5). Therefore, linking backward walking to pulse song specifically ensured a range of behavioral contexts (though always around song production) for learning. This allowed us to investigate the effect of perturbed social experience on male courtship, by optogenetically inducing a natural behavior—backward walking—in an atypical context—courtship.

**Fig. 1.**
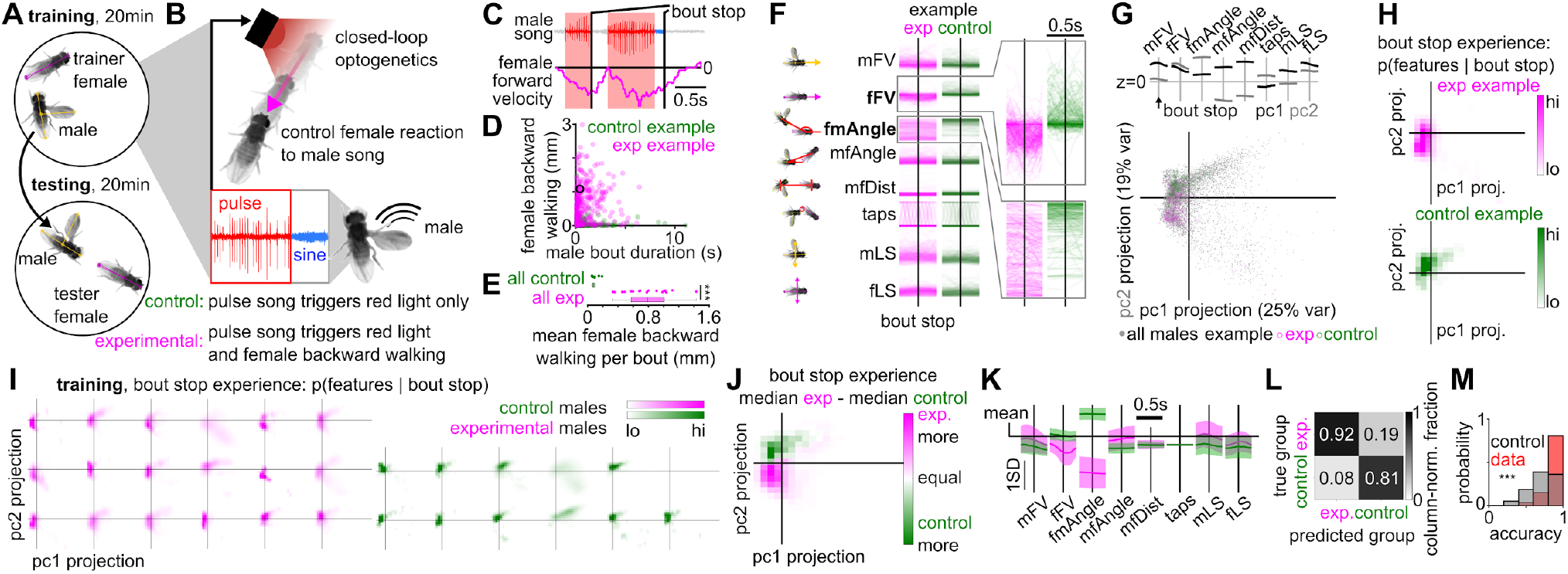
A novel assay for social experience-dependent plasticity of *Drosophila* behavior. (**A**) Experimental design. In a first ‘training’ experiment, wild-type males are paired with a trainer female providing perturbed social feedback to their courtship song. In a subsequent ‘testing’ experiment, trained males are paired with a female providing unperturbed feedback to their song. (**B**) To perturb female feedback to male song during training, male song drives female backward walking via optogenetic activation of Moonwalker Descending Neurons in the female (MDN-splitGal4; UAS-csChrimson). Control males undergo training experiencing unperturbed feedback to their song. (**C**) Example male song bouts and corresponding female forward velocity. Velocities below the zero line indicate backward walking. Vertical lines mark the end of individual song bouts. (**D**) During training, male bout duration reliably controls backward walking in experimental female flies (magenta), but not in controls (green). (**E**) Mean female backward walking per male song bout. Each point is the average for a single recording. (**F**) Behavioral features extracted from audio and video recordings around the time of male bout stops, for one experimental (magenta) and one control recording (green). Zoom-ins shown for female forward velocity (fFV) and female-to-male angle (fmAngle). (**G**) Dimensionality reduction (principal component analysis, PCA, followed by projection onto the first two PCs) performed on bout stop-centered behavioral features of all recordings (n=18 experimental, n=11 control). Colored points correspond to features shown in F. (**H**) Probabilistic representation of bout-stop experience for examples shown in F, derived from behavioral features embedded in PC space (see Methods). (**I**) Bout-stop experience for individual males during training (magenta for experimental, green for control group). (**J**) Difference in the median bout-stop experience between experimental and control groups during training. (**K**) Movement features corresponding to the largest positive (magenta) and negative (green) interconnected regions in J. These regions are associated with increased bout-stop probability during testing, in males with or without prior aberrant social experience, respectively. (**L**) Performance of a linear classifier (see Methods) in predicting experimental vs. control group identity from bout-stop experience during training, averaged over 1001 bootstrap repetitions. For each repetition, classifier training and testing was performed on random 80 and 20% samples of the data. (**M**) Classifier prediction accuracy (number of correct predictions /number of predictions) over 1001 bootstrap repetitions (‘data’, red), compared to the performance of a classifier that always predicts the majority class (‘control’, gray). For M: Wilcoxon rank-sum test for equal medians, ***P<0.001.

Our assay comprises two experiments (**Fig. 1A**): first, a 20-minute *training* of naive single virgin males paired with single virgin females providing perturbed (experimental) or unperturbed (control) feedback yoked to courtship pulse song, and a subsequent 20-minute *testing* of the trained males with a new, unperturbed (naive) single virgin female (**Fig. 1B**). During training for experimental males, female backward walking typically reaches its peak at the end of a song bout (**Fig. 1C-D**). Although the song of control males also triggers an optogenetic stimulus LED, it does not drive female backward walking, because control males are paired with females without an active opsin (**Fig. 1E**, see Methods).

Fly social communication is highly variable with males and females dynamically influencing each others’ behaviors (6, 19). Therefore, the effect of perturbing female feedback during male singing is not a priori known. We therefore used an unbiased method that incorporates a broad range of behavioral parameters to uncover which aspects of male experience differ between experimental and control conditions. Similar to previous approaches (25, 26), we derived male and female movement features from automated pose estimation (27) to form a high-dimensional representation of the social interaction, and projected these features into a low-dimensional representation (here using principal component analysis, PCA) to facilitate subsequent analysis.

To quantify the social experience of males around the time of singing (and specifically around the end of a song bout) during training, we extracted movement features (including male and female forward velocity (mFV and fFV) and lateral speed (mLS and fLS), male tapping (taps), male-female distance (mfDist), the angle of the male thorax relative to the female body axis (fmAngle) and the angle of the female thorax relative to the male body axis (mfAngle)) from automated SLEAP pose estimation data (27), and arranged the data into 0.5 second-long chunks around the end of male song bouts (**Fig. 1F**). We reduced the dimensionality of the resulting data (pooled across recordings) via projection onto the first two principal components (**Fig. 1G**; see Methods), and used the resulting embedding to define male experience around song bout ends (what we refer to as ‘bout-stop experience’) as the probability for a particular combination of movement features to occur around the time of song bout end – *p(features* | *bout stop)* (**Fig. 1H**). This definition of song experience ensured that the female reaction to male song was captured. Compared to controls, the bout-stop experience of experimental males (but not control males) was consistently characterized by female backward walking and situations in which the male was positioned closer to the head of the female (the latter being associated with small values of the female-to-male angle, fmAngle, **Fig. 1G-K**), such that a linear classifier trained on experiences of a subset of males was able to accurately predict the group identity of held-out song experiences (compared to a control classifier that predicts the majority class; **Fig. 1L-M**). These results confirm that optogenetic perturbation of MDN neurons in females alters the courtship singing experience of experimental males, which we quantify in more detail below.

### Characteristics of the Aberrant Social Experience

We next investigated whether female backward walking induces a consistent single bout-stop experience, or multiple discrete experiences not had by control males. Addressing this required, first, identifying the set of ‘base experiences’ that capture the variability between all males (both experimental and control), and second, determining the subset of base experiences associated with experimental males. Choosing a probabilistic description of the male’s bout-stop experience allowed us to use non-negative matrix factorization (NMF, see Methods (28)) to identify a set of eleven ‘base experiences’ that explains the variability of all bout-stop experiences during training. Briefly, NMF approximates each individual male’s bout-stop experience as a weighted sum of the identified base experiences (**Fig. 2A**), and the number of base experiences is chosen to minimize the Akaike Information Criterion (AIC (29), see Methods, **Fig. 2B**). Inspecting the weights of individual base experiences revealed three types of experiences: A first subset of experiences was overrepresented in control males, such as tapping the abdomen of a stationary female at short male-female distance and at intermediate to large female-to-male angles (base experiences 9-11 in **Fig. 2C**; yellow circles in each shown base experience mark the most likely feature combination given a bout stop, and these feature combinations are shown below each base experience). A second subset of experiences was shared by experimental and control males, such as orienting towards or pursuing a fast-locomoting female (base experiences 6 and 7, **Fig. 2C**). A third subset of experiences was overrepresented in, or even unique to experimental males (experiences 1-4), such as the female moving backwards and away from the male (base experience 3, **Fig. 2C**). These results quantify the multiple new experiences of experimental males. We next asked whether these altered experiences during training affect courtship behavior with naive females during subsequent testing.

**Fig. 2.**
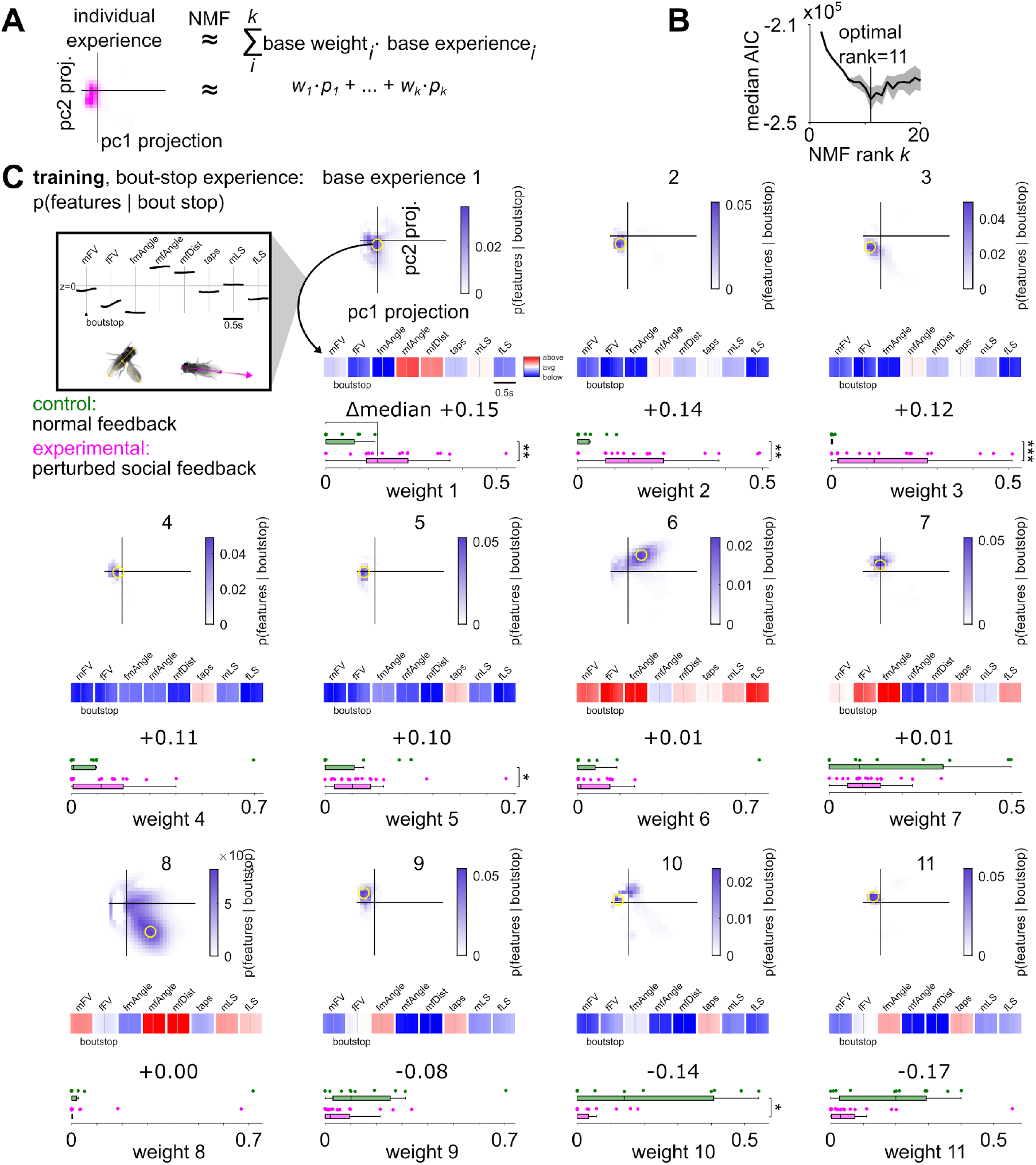
Male base bout-stop experiences during training. (**A**) Non-negative matrix factorization (NMF, see Methods) was used to approximate each individual male’s bout-stop experience (cf. **Fig. 1I**) as a weighted sum of base experiences. The number of base experiences, *k*, is a free parameter in NMF. (**B**) The number of NMF base experiences was chosen to minimize the median Akaike Information Criterion (AIC, median over 7 NMF runs with random initial conditions), yielding a set of *k*=11 base bout-stop experiences. (**C**) Automatically identified set of eleven base bout-stop experiences of (n=18 experimental and n =11 control) male flies during training. Yellow circles mark feature combinations that are most likely around the end of a song bout. Below each base experience, the most likely feature combination (pc1+pc2) at the time of ending a song bout is shown (corresponding to the yellow circle above). Below, the distribution of weights onto each base experience is shown for experimental and control males (magenta and green, respectively). Each individual male’s bout-stop experience is approximated by the weighted sum of base experiences, via non-negative matrix factorization (NMF; see Methods). Bout-stop experiences are shown sorted by the difference in median weight per group. Fly schematics illustrate the most likely movement feature combinations for base experience 1 and 11. For C: Wilcoxon rank-sum test for equal medians, ***P<0.001, **P<0.01, *P<0.05. The central mark indicates the median; the bottom and top edges of the box indicate the 25th and 75th percentiles, respectively.Whiskers extend to 1.5 times the interquartile range away from the box edges.

#### Males Alter their Song Usage Following Aberrant Social Experience

Thus far, we have focused on the experience of the male fly around the end of his song bouts, because the song-triggered modulation of female behavior peaks around this time during training (**Fig. 1C**). To determine whether this altered experience modulates the male’s strategy to initiate singing when paired with new females, we examined male-female dynamics at the start of a song bout. In principle, a change in strategy could comprise an overall change in the amount of song regardless of the male’s location relative to the female (as in learned suppression of song after experiencing female rejection (11)), or a change in song initiations linked to a specific social context (in other words, *when* and *where* to initiate singing, with respect to the female’s movements and position). To assess whether males showed either of these changes following aberrant social experience, we defined the male’s ‘bout-start strategy’ as the probability to initiate singing for any given combination of movement features – *p(bout start* | *features)*. Similar to our previous definition of bout-stop experience *p(features* | *bout stop)*, we extracted the same eight movement features from pose estimation data and arranged the data around the initiation of male song bouts. We reduced the dimensionality of the resulting data (pooled across recordings) via projection onto the first two principal components (see Methods), providing *p(features* | *bout start)*. We then projected the features of the entire recording (not only the instances of bout initiation) into the resulting embedding – providing *p(features)*. This allowed us to calculate *p(bout start* | *features)* – our measure of bout-start strategy – for each individual male.

We found that during testing, the bout-start strategy systematically differed between experimental and control groups (**Fig. 3A-D**). A linear classifier trained only on testing strategies (and hence blind to training data) predicted group identity with above-chance performance, supporting the conclusion that the perturbed social experience during training changed the males’ behavioral strategy used during testing (**Fig. 3F-G**). Overall, males with prior aberrant social experience were more likely than controls to initiate song when orienting towards a fast locomoting female at relatively large distances, whereas control males without aberrant social experience were more likely to initiate song when close to a stationary female (**Fig. 3E**), suggesting a social-context specific change in song strategy, rather than a global change in song probability. During training, song-triggered backward walking could be utilized by the male to stop a distant, fast-locomoting female, whereas singing towards a stationary female during training often leads to the male and female facing each other (indicated by small values of fmAngle and mfAngle, as for base experiences 2-5 in **Fig. 2C)**. The change in male song strategy observed during testing would be expected for a male that seeks to be close to the female abdomen, to be able to attempt copulation (typically while producing pulse song to trigger opening of the female vaginal plate (5, 30, 31)).

**Fig. 3.**
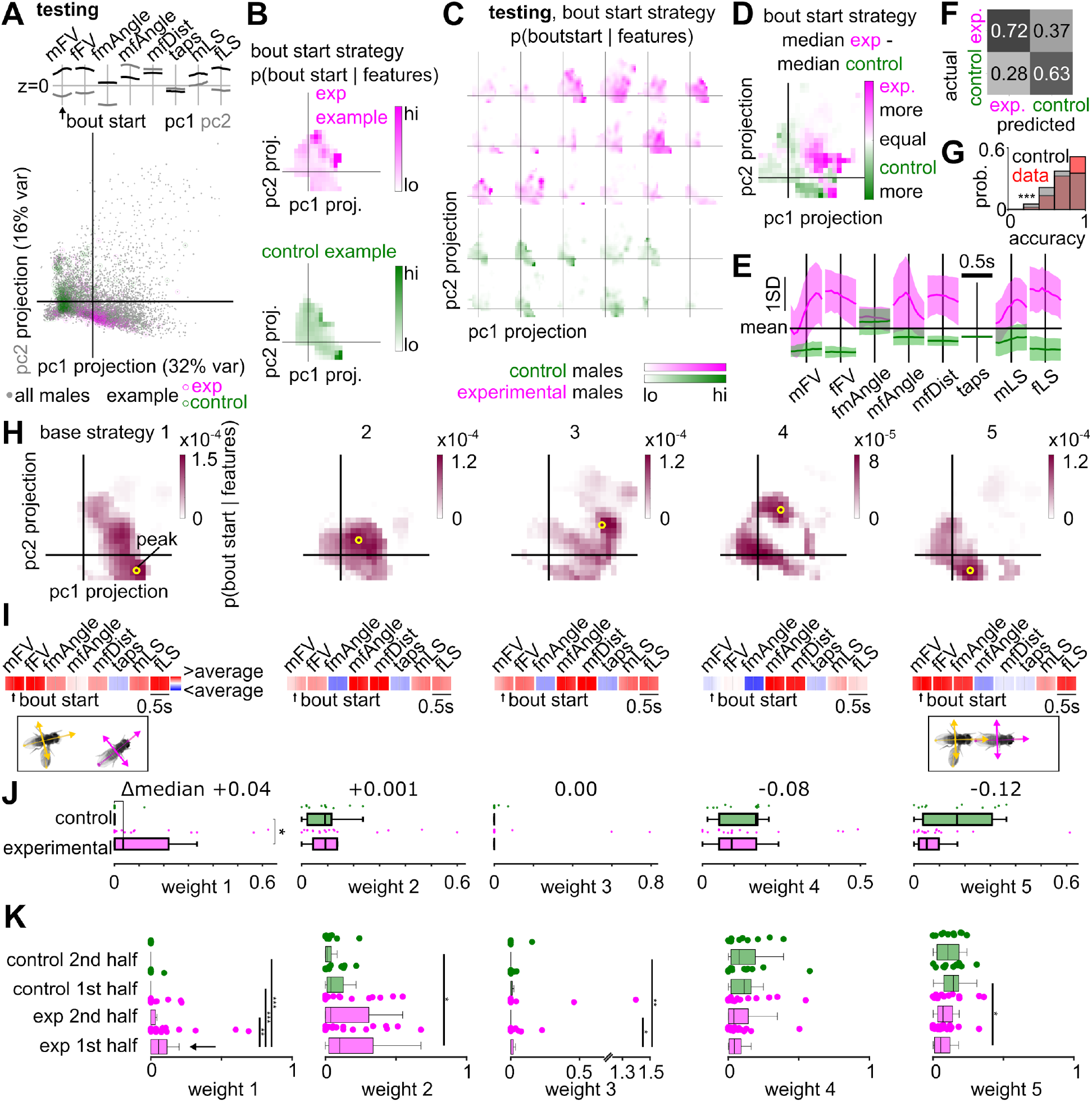
Unsupervised identification of base strategies used during testing. (**A**) Dimensionality reduction (principal component analysis, PCA, followed by projection onto the first two PCs) performed on bout start-centered behavioral features of all recordings (n=18 experimental, n=11 control). Colored points correspond to features from testing experiments for the males shown in **Fig. 1G**. (**B**) Probabilistic representation of bout-start strategy for examples shown in A, derived from behavioral features embedded in PC space (see Methods). (**C**) Bout-start strategy for all males of the experimental (magenta) and control group (green) during testing. (**D**) Difference of the median bout-start strategy between experimental and control groups during testing. (**E**) Movement features corresponding to the largest positive (magenta) and negative (green) interconnected regions in D. These regions are associated with increased bout-start probability during testing, in males with or without prior aberrant social experience, respectively. (**F**) Confusion matrix summarizing the performance of a linear classifier in predicting experimental vs. control group identity from bout-start strategies during testing. (**G**) Classifier prediction accuracy (number of correct predictions /number of predictions) over 1001 bootstrap repetitions (‘data’, red), compared to the performance of a classifier that always predicts the majority class (‘control’, gray). (**H**) Automatically identified set of five base bout-start strategies used by male flies during testing. Yellow circles mark feature combinations that maximize the probability to start a song bout. (**I**) Feature combinations (pc1+pc2) that maximize the probability to start a song bout (corresponding to the yellow circles in A). (**J**) Distribution of weights onto the base strategies in H for experimental and control males. Each male’s bout-start strategy is approximated by the weighted sum of base strategies (see Methods). Numbers at the top indicate the difference in median weight between groups; base strategies and weights in H-K are sorted in order of descending weight difference. (**K**) Bout-start strategy weights inferred for first and second half of song bouts in each recording (see Methods). Base strategy 1 is used by experimental flies mainly during the first half of testing bouts. For G, J-K: Wilcoxon rank-sum test for equal medians, ***P<0.001, **P<0.01, *P<0.05. The central mark indicates the median; the bottom and top edges of the box indicate the 25th and 75th percentiles, respectively.Whiskers extend to 1.5 times the interquartile range away from the box edges.

#### The Use of Novel Strategies Following Aberrant Social Feedback

Males could respond to perturbed social experience by forming novel behavioral strategies that are not part of the innate repertoire and hence missing in controls, or they could redistribute to what extent they use an existing set of ‘base strategies’. To investigate this, we again used NMF, now for unsupervised identification of the set of ‘base strategies’ that optimally represents *when* and *where* (relative to the female) the male sings during testing, such that the strategy of each male is approximated via NMF as a weighted linear sum of these base strategies (see Methods). Our analysis (**Fig. S2**) revealed a set of five base strategies, out of which four (base strategies 2-5) were used both by experimental and control males, but with distinct weight: strategies 2-4 were used to a similar extent by males with perturbed social experience and controls, whereas strategy 5 was primarily used by control males (**Fig. 3H-I**), suggesting a redistribution of strategy weights following perturbed social experience. In contrast, base strategy 1 was exclusively used by a subset of experimental males and not by controls, suggesting male flies can develop novel strategies through learning from past social experience.

#### What do the base strategies represent?

Overall, males with aberrant social experience traded strategy 5 for strategy 1. While both strategies corresponded to initiating song during fast male and female locomotion, with the male behind the female and oriented towards her, strategy 1 involved initiating song at larger male-female distance compared to strategy 5 (**Fig. 3H-I**). Notably, inducing female backward walking when the male is facing the female abdomen and moving at fast speed but at large distance is the context during which the backward walking brings the female abdomen closer to the male, and hence to a more favorable location to attempt copulation. Learning this would be advantageous for the male during training, but not during testing, since the male cannot induce backward walking during testing.

#### Males that experienced a backward walking female revert to default strategies after experiencing normal female behavior

Our results raise the question of whether the novel strategies observed for males with perturbed experience during training persist during ongoing experience with a normal female during testing. To test this, we compared the relative weight onto each base strategy during testing when using only the first vs. second half (during the courtship encounter with the test female) of song bouts for each male in the given NMF model. We found that usage of strategy 1 increased following aberrant social experience only for the first half of song bouts (**Fig. 3K**; the first half of bouts was produced largely during the first half of the recording, over a period of around 10 min on average, **Fig. S3A-B**), supporting the conclusion that in our assay, ongoing experience with a normal female can revert the use of novel strategies that were acquired following perturbed social experience. Whether repeated training, or the introduction of a consolidation period between training and testing can fortify novel strategies, as observed for courtship conditioning (11), remains to be tested.

## Discussion

In this study, we establish a novel assay for experience-dependent plasticity of courtship behavior in male *Drosophila melanogaster*. Our assay does not simply modulate the gain of behavior (producing more or less of a given courtship action, similar to social-experience dependent modulation of male courtship drive, or of aggression, (11, 32, 33)) but the strategy underlying behavior (when and where the behavior is produced in the social context). We find that experiencing perturbed social feedback from a female alters male courtship behavior towards subsequently paired females. Specifically, males that experienced female backward walking in response to their own courtship song change the conditions under which they produce song - most notably, they increase song production during chasing at above-average relative distances (control males also sing during chasing (although to a lesser degree), but at closer distances). Such experience-dependent altering of the context in which a behavior is executed has been found in other species (16, 17, 18), and is commonly referred to as ‘usage learning’.

### Learning Mechanisms and Timescales

Prior work in *Drosophila* has identified several genes (e.g., *dunce, rutabaga*) and neural cell types (e.g., dopaminergic) types involved in both social and non-social learning ((13); reviewed in (34, 35, 36, 37)). Further, for male flies during courtship, prior copulations increase the likelihood of abandoning an ongoing copulation in response to stress; this experience-dependent plasticity of mating behavior is mediated by dopaminergic signaling, as is male mating drive and courtship itself (38, 39, 40). Whether these mechanisms also contribute to the experience-dependent modulation of courtship strategies we describe remains to be tested. Given the timescales of learning and memory we report here (learning occurs over a 20 min period, and plasticity of the behavior is lost within, on average, 10 min with a new female), we postulate that dopaminergic modulation also plays a role, as dopamine-mediated plasticity has been shown to operate on timescales of several minutes (41, 42, 43). Further, the usage decay for the learned strategy observed during testing with a normally behaving female would be consistent either with memory decay as known from classical conditioning experiments in *Drosophila* (44) or extinction learning due to the lack of a reinforcing signal (here: female backward walking) during testing (45).

### Valence of the Aberrant Social Experience

What and how does the male learn from a backward walking female? And specifically, is a backward-walking female a positive or negative reinforcer to the male? Recent work has shown that artificial activation of Moonwalker Descending Neurons – as performed in our study – is aversive to the stimulated fly (46), suggesting that females could provide a negative reinforcement signal to the male during song-triggered backward walking (via, for example, the release of stress-related odorants (47)). On the other hand, courtship itself is appetitive to the male (40), suggesting that in our experiments, the male could be exposed to a mixture of appetitive and aversive signals. Specifically, the dominant valence could depend on social context, since backward walking can bring the female closer to the male (potentially appetitive) when she is far away and male and female orientation aligned, or send her away from the male (potentially aversive) when the male is lateral or frontal to the female when initiating song. Consistent with this hypothesis, the males in our dataset do not stop courting the female during training, as would be expected for an overall aversive stimulus such as ongoing female rejection in courtship conditioning (11), suggesting that potential backward-walking related aversive signals are sufficiently counterbalanced by courtship-activity induced appetitive signals in order to maintain male courtship drive.

### Neural Circuit Modulation

Learned suppression of a given behavior (as in courtship conditioning, see above) can be implemented via gain control of command neurons driving the behavior. In contrast, learned changes of a behavioral strategy beyond simple gain modulation likely requires more complex mechanisms. In the present work, we observed an experience-dependent shift to singing more in a specific behavioral context – when the female is far and moving fast, in front of the male. This could be due to modulation of activity in visual neurons encoding female motion during courtship (21, 22) or to modulation of song-driving downstream targets such as pC2l or pIP10 (5, 48, 49, 50). These hypotheses could be tested by recording from these cell types during learning in a courtship virtual reality.

### Prior work using closed-loop stimulation to create alternate realities and study learning

Closed-loop stimulation approaches have been used successfully to alter song production and learning, and to identify dopaminergic song-performance error signals, in birds (51, 52, 53, 54, 55) or to induce learned female avoidance in male flies (56). Our approach is novel in using an indirect modulation of the male’s social experience, via optogenetic perturbation of the female’s response to male song behavior. While we used female backward walking as an induced response to male song, our approach can be easily extended to induce any other behavior for which genetic driver lines are known. For example, by triggering female rejection in response to male advances (27). Future work can hence investigate which feedback cues can serve to modify male courtship behavior, and similarly, whether female behavior is also plastic.

### Generality and limitations of our method

Our analytical framework for quantifying experience-dependent plasticity of social strategies can be easily ported to any other system in which automated pose estimation is feasible. One limitation of our approach is its definition of the male song strategy based on the likelihood of initiating song given a particular movement feature combination. Estimation accuracy for this likelihood increases with the size of the sampling window, and therefore our approach has limited use for short recording durations.

To conclude, we used a novel assay to show that the innate courtship strategy of male *Drosophila* is shaped by past social experience via usage learning, demonstrating behavioral plasticity beyond simple gain modulation. Given the wealth of genetic and neural circuit tools available for this model system, including recently released wiring diagrams of the whole brain and ventral nerve cord (57, 58, 59), it should be possible to discover the precise circuit mechanisms that shape usage learning.

## Materials and Methods

### Experimental design

Behavioral experiments were performed in two custom made circular chambers (modified from (4), as described in (5)) within black acrylic enclosures. Ambient light was provided through an LED pad inside each enclosure (3.5” × 6” white, Metaphase Technologies). For each chamber, video was recorded at 60 fps (FLIR Blackfly S Mono 1.3 MP USB3 Vision ON Semi PYTHON 1300, BFS-U3-13Y3M-C, with TechSpec 25mm C Series VIS-NIR Fixed Focal Length Lens) using hardware triggering and using infrared illumination of around 22 μW/mm2 (Advanced Illumination High Performance Bright Field Ring Light, 6.0” O.D.,Wash Down, IR LEDs, iC2, flying leads) and an infrared bandpass filter to block the red light used for optogenetics (Thorlabs Premium Bandpass Filter, diameter 25 mm, CWL = 850 nm, FWHM = 10 nm). Sound was recorded at 10 kHz from 16 particle velocity microphones (Knowles NR-23158-000) tiling the floor of each chamber. Microphones were hand-painted with IR absorbing dye to limit reflection artifacts in recorded videos (Epolin Spectre 160). Temperature was monitored inside each chamber using an analog thermosensor (Adafruit TMP36). Before each experiment, one chamber was randomly chosen to drive closed-loop stimulation. Sound recorded in the closed-loop chamber was analyzed in real time to detect male pulse song. Red LEDs for optogenetic activation in both chambers were turned on upon detection of pulse song in the closed loop chamber, and turned off if no pulse was detected. At the beginning of each experiment, flies were gently aspirated into the chambers, and the experiments were timed to start within 120 minutes of the incubator lights turning on, to use flies during their first activity peak. Flies first performed a 20 minute training experiment, after which both flies were gently extracted from each chamber. Males were reinserted into the chamber with a naive virgin female to perform a 20 minute testing experiment.

### Data exclusion criteria

Only recordings with more than or equal to 25 pulse trains were kept for analysis, to ensure sufficient data was available to estimate conditional probability densities (song experience and song strategy).

### Prevention of copulation

To reduce the rate of copulation and hence ensure equal training time for all male flies, the male labellum was removed under cold anesthesia 30 minutes prior to an experiment. This procedure proved effective to minimize copulation rate while maintaining male courtship vigor.

### Optogenetics

The red-shifted channelrhodopsin CsChrimson (23) was expressed in Moonwalker Descending Neurons (MDN1) of female flies (VT44845@attP40 ZpGAL4DBD, VT50660@attP2 p65ADZp (24)). Flies were kept for 3-5 days on regular fly food or food supplemented with all-trans retinal (ATR, cofactor required for functional expression) at 1 ml ATR solution (100 mM in 95% ethanol) per 100 ml of food. CsChrimson was activated at 50μW/mm2, using 627nm LEDs (Luxeon Star). The duration and pattern of stimulation was determined by the pulse song dynamics of a given male fly.

### Fast online song segmentation

To enable closed-loop optogenetic stimulation triggered on the detection of male pulse song, fast online song segmentation was performed as described previously (5). The data acquisition (DAQ) device delivered 8 ms of 16 channel sound per read out, which we combined with 16.5 ms of history to have 24.5 ms of sound for pulse detection, exceeding the duration of a typical pulse. Both for building a data set for the classifier during training and for preparing a window of audio recording collected in real time for the classifier during real-time classification, the raw waveform was preprocessed as follows. For every 245-sample (24.5-ms) window, the channel with the maximum-power sample was selected for further processing. The single-channel window was then normalized: The window was first divided by its 2-norm and then smoothed using moving-window averaging with a 15-sample filter. The center of the candidate pulse was identified as the sample with the maximum power within the central 80 samples in the smoothed window. The window was then cropped to the 165 samples centered on the candidate pulse. The amplitudes of the candidate-pulse waveform were inverted if the 10 samples preceding the center had a negative mean. This ensured that no polarity reversals due to recording artifacts impaired the training or the classification procedure. The normalized 165-sample waveform served as input to a convolutional neural network (CNN). In the first layer, eight filters of 9-sample width were convolved with the input using a stride length of 1 sample. The layer was activated using a rectifier linear unit (ReLU). In the second layer, eight filters of 9-sample width were convolved with stride length of 2 samples. Upon ReLU activation, the outputs were max-pooled with pooling width of 2 samples. The resulting outputs were flattened, and put through a ReLU-activated dense network, resulting in a layer of 32 neurons. The 32 neurons were put through a sigmoid-activated dense network resulting in one neuron, whose value between 0 and 1 was correlated with the probability that the given waveform had a pulse in the center. 25%-dropouts were applied after the flattening stage before the sigmoid activation to prevent overfitting. For each audio input, the CNN produced a value between 0 and 1. The closer this value was to 1, the more likely it was that the song window contained a pulse in its center. We considered output values exceeding 0.99 a pulse. Any pulse candidates separated by IPIs less than the 15ms from the previously detected pulse were ignored. For interfacing the DAQ device with the Python software developed for song mode detection and response, PyDAQmx was used (https://pythonhosted.org/PyDAQmx/). The CNN was implemented using Keras with Theano backend (https://github.com/keras-team/keras, (60)). 19 song recordings from behavioral assays for unique NM91 (wild-type) male-female pairs were used to generate the data sets used for training, validation, and evaluation of the CNN. Windows of songs were labeled as pulse/no-pulse, using outputs from the offline song segmenter as the ground truth. For the no-pulse class, the windows contained no regions of pulse. For the pulse class, the windows included one pulse. 11, 4, and 4 recordings were used for training, validation, and evaluation, respectively. During training, class weights were applied such that the less represented class would have, on average, equal loss as the overrepresented class in a completely naive, untrained classifier.

### Offline song segmentation

For subsequent offline analysis, song was segmented as described previously (4), using a modified sine detection parameter to account for different acoustics in the modified setup used here (Params.pval = 1e-7).

### Animal tracking and pose estimation

Male and female pose (locations of head, thorax, left and right wing tip) was automatically estimated and tracked, and manually proofread for all videos using SLEAP (27).

### Choice of movement features

From pose estimation data, a total of eight movement features were calculated: male and female forward velocity (mFV, fFV), male and female lateral speed (mLS, fLS), male taps (binary variable, indicating whether a male did or did not tap the female abdomen in a given video frame, as described in (5)), as well as relational features male-female distance (mfDist), female-to-male angle (fmAngle, 0° when the male is in the front of the female, 180° if he is behind the female), and male-to-female angle (mfAngle, 0° when the female is in front of the male, 180° if she is behind the male).

### Behavioral readouts of social experience and behavioral strategies

Movement features characterizing male-female interactions (see previous paragraph) were extracted from pose estimation data, and for each recording organized into overlapping chunks of 5000 samples (0.5s before until after each point). Chunks containing a song event (start or end of a pulse train, depending on whether the recording was from a testing or training experiment) were pooled across recordings of both the experimental and control group, and principal component analysis followed by projection onto the first two principal components (PCs), of the song-triggered features was used to define an ‘experience space’ that all feature chunks (including those not containing song) were projected into. Each point in this space corresponds to a particular social experience characterized by the projection onto the first two PCs. The probability density of all song-triggered chunks of one male defined his ‘song experience’, *p(feature combination* | *song event)*, whereas the probability of all chunks of that male defined his ‘overall experience’, *p(feature combination)*. Similarly, male ‘song strategy’ was defined as the probability to sing given the PC projection of any feature combination, *p(song event* | *feature combination)*.

### Movement features for overrepresented experiences and strategies

Overrepresented experiences and strategies were defined as regions in behavior space for which the difference between the group-level median bout-stop experience (training) and bout-start strategy (testing) was larger (experimental group) or smaller than zero (control group), respectively. These regions of parameter space were binarized, and connected components analysis was used on the resulting binary maps to identify congruent overrepresented regions. For the largest connected component, the movement features were identified.

### Unsupervised identification of base experiences and base strategies based on non-negative matrix factorization

Defining social experience and behavioral strategy as probability densities (that is, non-negative and normalized; see above) allowed us to use non-negative matrix factorization (NMF, (28)) to approximate each male’s experience and strategy as a weighted sum of ‘base experiences’ and ‘base strategies’. The number of such base components, *k*, is a free parameter in NMF, and how well NMF approximates the data depends on the choice of *k*. We used the Akaike Information Criterion (AIC (29)) to automatically identify the optimal *k* that minimized:

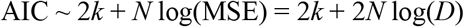

where *N* is the number of data points (elements per experience or strategy × number of recordings), MSE is the mean squared error of the NMF reconstruction, and *D* is the root mean squared error (as returned by Matlab’s nnmf.m function). Specifically, we ran 7 NMF runs with random initial conditions and chose the *k* that minimized the median AIC over all seven runs. While the above form of AIC assumes Gaussian residuals, we use it as a model selection heuristic. Since the same residual structure applies across values of *k*, differences in AIC remain informative for comparing models, even if the exact noise distribution is unknown.

### Inference of NMF weights for partial recordings

To infer base-strategy weights using only the first or second half of song bouts in each recording, we calculated each male’s bout-start strategy using the first or second half of song bouts, and then ran the NMF algorithm, fixing the base strategies as those inferred for all song bouts.

### Linear discriminant analysis of group separability

To quantify how well experimental and control groups were distinguished based on individual experiences or strategies, we trained linear classifiers on 80% of a given dataset and used the trained classifiers to predict group identity on the held-out 20%. This procedure was repeated 1001 times using random samples, confusion matrices were calculated per repetition and averaged across repetitions to provide a summary statistic. Prediction accuracy (number of correctly predicted labels /number of labels) was also calculated per repetition, and compared to the accuracy of a classifier that always predicts the majority class. Confusion matrices for testing experiments (predicting group identity based on the bout-start strategy used during testing) were used to assess the strength of experience-dependent behavioral plasticity.

### Genotypes

See **Table 1** for a list of genotypes used in this study.

**Table 1.** Key resources and reagents.

| Reagent type (species) or resource | Designation | Source or reference | Identifiers | Additional information |
| --- | --- | --- | --- | --- |
| Genetic reagent (*D. mel*) | NM91 | (6) |  | Obtained from Peter Andolfatto |
| Genetic reagent (*D. mel*) | VT44845-ZpGAL4DBD (attP40) | (24) | BDSC: 72509 | Obtained from Barry Dickson |
| Genetic reagent (*D. mel*) | VT50660-p65ADZp (attP2) | (24) |  | Obtained from Barry Dickson |
| Genetic reagent (*D. mel*) | 20xUAS-IVS-CsChrimson. mVenus (X) (attP18) | (23) | BDSC: 55134 | Obtained from Vivek Jayaraman and Gerry Rubin |

### Declaration of generative AI and AI-assisted technologies in the writing process

During the preparation of this work the authors used GPT-4o in order to improve language and readability. After using this tool, the authors reviewed and edited the content as needed and take full responsibility for the content of the publication.

## Funding

HHMI Faculty Scholar award (MM)

NIH NINDS Research Program R35 award (MM)

NIH BRAIN R01 (MM)

German Research Foundation grant RO 5787/1-1 (FAR)

German Research Foundation grant RO 5787/2-1 (FAR)

Klaus Tschira Boost Fund, a joint initiative of the German Scholars Organization and the Klaus Tschira Foundation, grant GSO/KT59 (FAR)

zukunft.niedersachsen program of the Ministry for Science and Culture Lower Saxony, grant 15-76251-2/24-4058/2024 (FAR)

Burroughs Wellcome Fund PDEP and the SCGB BTI Award (OMA)

## Author contributions

Conceptualization: FAR, MM

Methodology: FAR, AJC, MC, OMA, MM

Investigation: FAR, MM

Data analysis: FAR

Visualization: FAR

Funding acquisition: FAR,

MM Supervision: MM

Writing – original draft: FAR, MM

Writing – review & editing: FAR, ECI, AJC, MC, OMA, MM

## Competing interests

The authors declare no competing interests.

## Data and materials availability

All data needed to evaluate the conclusions in the paper are present in the paper and/or the Supplementary Materials.

## Notes

### Competing Interest Statement

The authors have declared no competing interest.

