## Supplemental Information for "Recent social experience alters song behavior in *Drosophila*"

Frederic A. Roemschied *et al.*

**This pdf includes:**

Figs S1 to S3

**Fig. S1.**

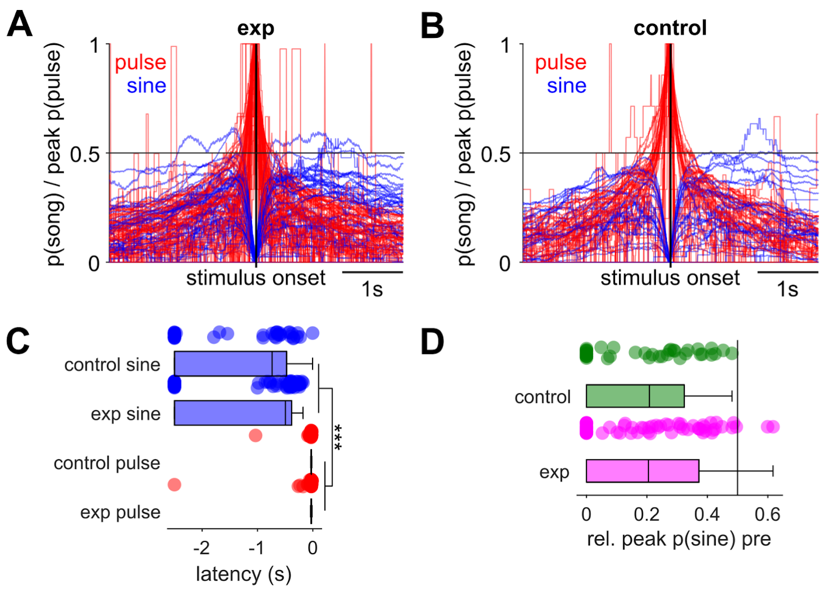


**Feedback perturbation is specific to pulse song.** (**A**) Probability for n=18 experimental male flies during training to sing pule (red) or sine song (blue) as function of time from optogenetic stimulus onset, and normalized by the peak pulse probability for each male. (**B**) Same as A but for n=11 control males. (**C**) Latency from peak pulse (red) or sine song (blue) to stimulus onset, considering only the time preceding stimulus onset. Pulse song consistently preceded stimulation with short latencies (median latencies 28.3ms and 30ms for experimental and control males). (**D**) Peak probability of sine song preceding stimulation, relative to peak pulse song probability preceding stimulation. The peak sine probability preceding stimulation consistently amounted less than 50% of the peak pulse probability. (median 20.8% and 20.5% for experimental and control males, respectively). For C-D: Wilcoxon rank-sum test for equal medians, ***P<0.001. The central mark indicates the median; the bottom and top edges of the box indicate the 25th and 75th percentiles, respectively. Whiskers extend to 1.5 times the interquartile range away from the box edges.

**Fig. S2.**

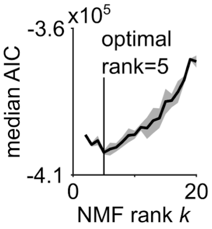


**Optimal number of base bout-start strategies during testing.** The number of NMF base bout-start strategies was chosen to minimize the median Akaike Information Criterion (AIC, median over 7 NMF runs with random initial conditions), yielding a set of *k*=5 base bout-start strategies.

**Fig. S3.**

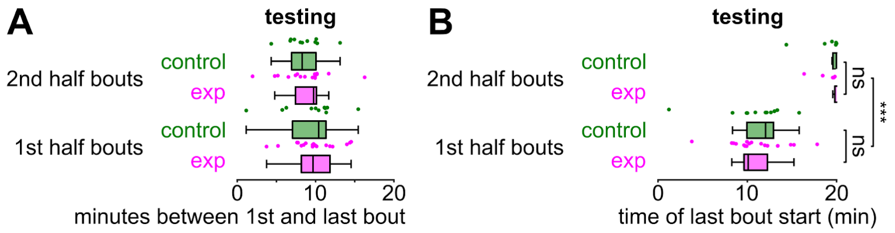


**Period over, and time at, which first and second half of song bouts were produced during testing.** (**A**) The first and second half of song bouts were produced over a period of around 10min on average, with no significant difference between the medians of the experimental and control groups (nor first and second half). (**B**) For the first and second half of song bouts, the respective last bout was initiated around 10-12 and 20 minutes into the testing experiment, respectively. A-B: Wilcoxon rank-sum test for equal medians; the central mark indicates the median; the bottom and top edges of the box indicate the 25th and 75th percentiles, respectively.Whiskers extend to 1.5 times the interquartile range away from the box edges.
